# A novel mechanism for centrosome expulsion ensures metabolic activity in polyploid cells

**DOI:** 10.1101/2025.08.01.668089

**Authors:** Riham Salame, Simon Gemble, Anthony Simon, Lin Jawish, Anne-Sophie Mace, Ptissam Bergman, Graca Raposo, Clotilde Cadart, Renata Basto

## Abstract

Programmed polyploidy is often linked to increase cell size to support enhanced metabolism, barrier function or regeneration. In certain polyploid cells, the cytoskeleton is drastically remodeled-most notably by eliminating centrosomes. However, the purpose and mechanisms underlying centrosome elimination have remained unclear. We investigated this question in *Drosophila* acentrosomal salivary glands (SGs), a physiological polyploid model where cells reach high chromosome content through endoreplication. Using genetic tools, live imaging approaches, super-resolution microscopy combined with tissue clearing and electron microscopy, our study uncovers a novel centrosome elimination pathway *in vivo*. This process requires non-muscle myosin II (MyoII) activity and the macroautophagic machinery to drive centrosome release into the lumen of the salivary glands via autolysosomal exocytosis. Failure to eliminate centrosomes disrupts the mitochondria network, impairing respiration and ATP production. Our findings reveal a previously unknown mechanism that removes centrosomes through the secretory autophagy pathway to protect mitochondrial function and support the high metabolic demands of polyploid cells.

## Introduction

Centrosomes are the main microtubule (MTs) organizing centers (MTOC) of animal cells, with key roles in polarity establishment, cell migration and cell division (*1*). Each centrosome contains two centrioles that recruit and organize the pericentriolar material (PCM), which is the site for MT nucleation (*2*, *3*). Most cycling cells maintain stable centrosomes numbers through the tight control of the centriole duplication machinery along the cell cycle (*4*, *5*). Moreover, surveillance mechanisms discard cells with abnormal centrosome number. Differentiated cells, such as neurons keep centrosomes, even if MTs are nucleated from alternative MTOCs, whereas multiciliated cells, massively amplify centrioles to be able to grow multiple cilia (*6–8*). In most animal oocytes, centrosomes are eliminated before fertilization (*9–13*). All these examples represent cells with diploid or haploid genome content that remain arrested in G0 or G1.

Polyploid cells contain more than two sets of all chromosomes. Physiological polyploidy is associated with cell size increase and increase in metabolic capacity, barrier function or regeneration (*14*, *15*). Interestingly, many polyploid cells generated through endoreplication, such as *Drosophila* salivary glands, ring gland cells or cells from the fat body do not contain centrosomes (*12*, *16*, *17*). This is also the case of *Drosophil*a and mammalian muscle cells, which are multinucleated, but in this case formed through cell fusion (*12*, *18*). Why and how certain polyploid cells lose their centrosomes is unknow. Here we investigated the mechanisms involved in centrosome elimination in *Drosophila* salivary gland (SG) polyploid cells. These are highly secretory tissues, which undergo up to 10 rounds of endoreplication generating cells with 1024C-2048C (C-value representing the DNA content in the haploid genome). In these cells, multiple DNA strands remain attached forming polytene chromosomes (*19–21*). Here, we found that during embryogenesis, centrosomes are released into the lumen of the salivary glands in a process dependent on non-muscle myosin II (MyoII) activity and autolysosomal exocytosis, to be subsequently targeted for degradation by the proteasome. Strikingly, our work shows that if centrosomes are maintained in SG cells, the mitochondrial network appears fragmented, and cells display reduced oxygen consumption rates. This study thus provides a direct link between centrosome elimination in programmed polyploid cells and metabolic processes essential for their function and homeostasis.

## Results

### Centrosomes are released into the salivary gland lumen during embryonic development

To investigate the steps involved in centrosome elimination in polyploid cells from *Drosophila* salivary glands (SGs), we analyzed embryonic development after stage 10, when SG cells are specified (*22*, *23*). In crowded wild type (WT) embryos, SGs were identified using an endogenous GFP tagged version of the transcription factor fork-head (Fkh) (*24–26*). Using antibodies that recognize the centriole and PCM proteins, Sas-4 and Pericentrin-like protein (Plp), respectively (Figure 1A-D, Supplementary Figure 1A), two centrosomes per cell were seen (Figure 1A and 1E) at stage 10, when the SG placode is formed (*22*, *23*, *27–29*). At stage 11-12, when SG cells start to invaginate, due to apical constriction induced by actomyosin contraction (*22*, *23*, *30*), approximately 50% of the centrosomes were localized in close proximity to the apical membrane (Figure 1B and 1E). During stages 13-14, the SG secretory cells start to posteriorly elongate, forming a tube-like structure with a lumen (*22*, *30*). At this stage, we noticed that certain centrosomes were located extracellularly, within the lumen of the salivary glands (Figure 1C and 1E) and at the later stages 15-17, the frequency of centrosomes in the lumen increased (Figure 1D-E).

**Figure 1.**
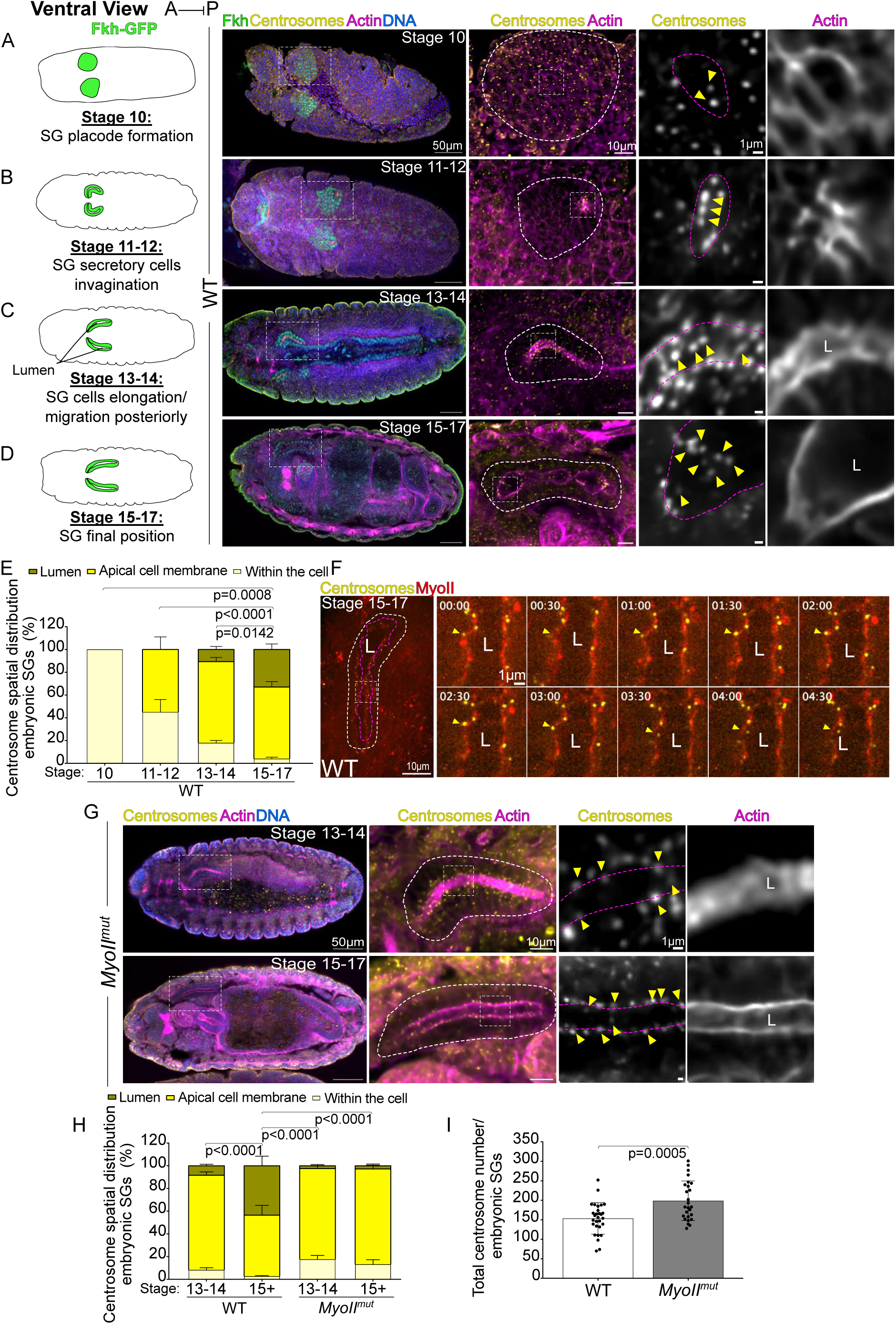
Centrosomes are transported towards the SG lumen during embryonic development. (A-D) On the left, drawing depicting the indicated embryonic stages according to Fkh expression to label SGs. On the right, representative immunostaining images showing low magnification views of embryos (left hand side) and higher magnification views (middle panels). Fkh in green, centrosomes in yellow, actin in magenta and DNA in blue. The white lines surround SGs, and the squares surround the region shown on the right-hand side, for centrosomes (left) and actin (right) depicted as separate panels in grey. In all insets, the yellow arrowheads point at centrosomes. In (A), the pink dashed line surrounds an SG cell. In (B), the centrosomes closely associated with the apical membrane during apical constriction. In (C-D) the pink dashed lines mark the lumen (L). (E) Graph bar showing the centrosome spatial distribution in SGs across different developmental stages as indicated. (F) Stills of time lapse movie of embryonic SGs expressing Sas4-GFP to follow centrosomes (yellow) and Myosin II (MyoII)-mCherry transgene (red) at stage 15-17. On the left an overview of the entire SG. The white and magenta dashed line surrounds the SG and the lumen (L), respectively. The square indicates the region shown at higher magnification stills on the right. The yellow arrowheads points to the trajectory of a centrosome that is initially located at the apical membrane and that is subsequently translocated into the lumen. (G) On the left, representative images of *MyoII^mut^* embryos of the indicated embryonic stages showing centrosomes (yellow), actin (magenta) and DNA (blue). In the middle panel, centrosomes and actin are shown. The white dashes lines surround SGs, and the white squares mark the regions shown on the righthand insets for centrosomes (top) and actin (bottom) depicted as separate panels in grey. The yellow arrowheads indicate centrosomes and the pink dashed lines the lumen. (H) Bar and dot plot graph showing the centrosome spatial distribution in SGs in WT and *MyoII^mut^* embryos at the indicated developmental stages. (I) Bar and dot plot graph showing the total number of centrosomes in SGs of the indicated genotypes. Bars show the mean ± SD. Statistical significance is shown and it was determined by two-way ANOVA tests (E, H) and unpaired t-test (twotailed) (I). In (E and H,) P value shown for the category of centrosomes in the lumen. For each graph at least three experiments were quantified and a minimum of 10 embryos/stage (E), minimum of 13 embryos/stage (H) and minimum 26 embryos/stage (I)

These observations suggest that centrosomes translocate from the cell to the lumen of the SG, in a process that remains to be understood. Using time-lapse microscopy, we followed the dynamics of centrosomes in embryos expressing Sas-4-GFP (*27*) and the regulatory light chain of non-muscle myosin II (NMMyoII) - Spaghetti squash (*31*, *32*) fused to mCherry (MyoII-mCh) to label cell boundaries and the lumen border. Strikingly, centrosomes were seen lingering around the apical membrane in an erratic movement before being transported into the lumen (Figure 1F and Supplementary movie1).

The MyoII-mCh signals were relatively low and due to photobleaching, we could only film for short periods of time. We also followed actin behavior using the LifeActin-RFP (*33*) reporter line, but the combination with Sas-4-GFP was not viable. We therefore labelled fixed embryos with phalloidin to label actin and phospho-myosin (p-MyoII) antibodies, which reveal active MyoII. Interestingly, in stage 15 embryos, actin positive vesicles were noticed positioned at the apical membrane and frequently these contained centrosomes. Analysis of actin vesicles containing centrosomes showed that 52% of these were also p-MyoII positive (Supplementary Figure 1B). The number of actin vesicles containing centrosomes was very variable and infrequent at the single embryo level, suggesting that this is a rapid event. Taking together, these results show that centrosomes are transported from within SG cells to the lumen, in an expulsion trajectory using vesicles enriched with actin and active MyoII.

### Centrosome expulsion depends on MyoII activity

To functionally test whether MyoII is involved in centrosome release into the SG lumen, we used a hypomorphic mutation in *spaghetti squash* (*sqh^mut^,* here referred to as *MyoII^mut^*) (*31*). Strikingly, analysis of *MyoII^mut^*embryos from stage 15 onwards revealed that centrosomes were mainly present in SG cells and only few centrosomes were detected in the lumen (Figure 1G-H). Using three-dimensional (3D) reconstructions, the difference in luminal centrosome number was obvious between WT and *MyoII^mut^* (Supplementary movies 2-3). Further, analysis of *MyoII^mut^* SGs, showed that centrosomes were localized within the SG cell, but close to the apical membrane (Figure 1H), suggesting that their positioning at this location is independent of MyoII.

Unexpectedly, careful analysis of *MyoII^mut^* SGs showed an increase in total centrosome number, when compared to WT SGs (Figure 1I). These observations, suggest that when centrosomes do not undergo their programmed elimination, through expulsion towards the lumen, they continue to duplicate throughout embryonic SG development, leading to centrosome amplification.

A prediction from the findings described above is that *MyoII^mut^* SG cells at later developmental stages should contain centrosomes, while WT SGs should not. We therefore analyzed both WT and *MyoII^mut^* SGs from third instar larval stages (L3) (Figure 2A and Supplementary Figure 2A), labelled with different centrosome markers (Supplementary Figure 1D-F). As expected, in WT SGs, centrosomes were absent from SG cells, while they could still be noticed in the lumen (Figures 2B-C, Supplementary Figures 2B-D). In contrast, in *MyoII^mut^* SGs, multiple centrosomes were readily identified in the large majority of SG cells (Figures 2B-C), while the lumen only occasionally contained centrosomes (Supplementary Figures 2B-C). We confirmed that the centrosomes present in SG cells, contained the expected centriolar structure (Figure 2D) as previously described when using optically demodulated structured illumination techniques (SIM) (*34*).

**Figure 2.**
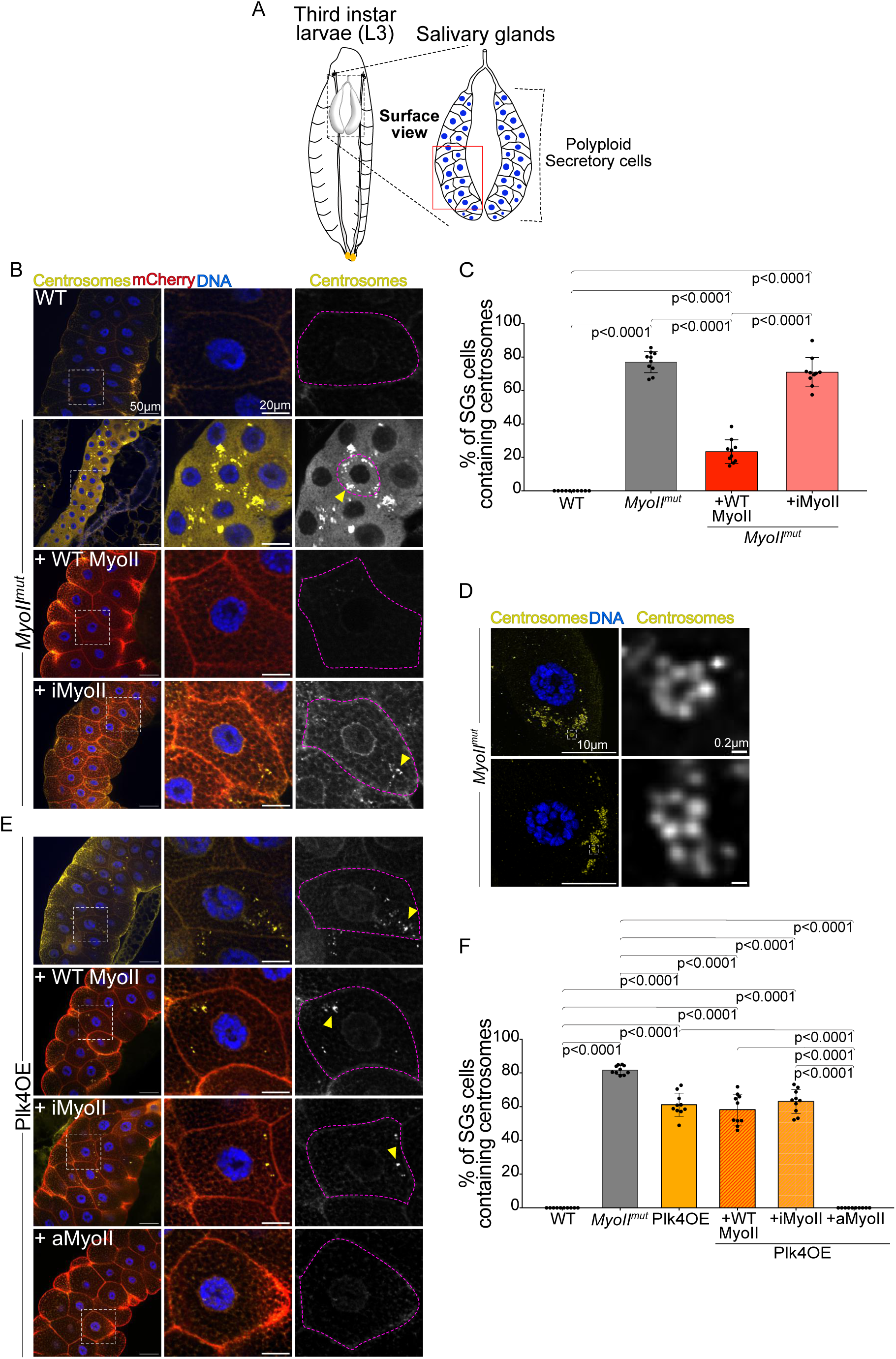
Centrosome expulsion requires MyoII activity. (A) Schematic drawing of SGs from third instar larvae (L3). On the left an L3 larvae is depicted with anterior positioned at the top. On the right, the SG pair showing a top/surface view of polyploid secretory cells with nuclei depicted in blue. (B and E) On the left, representative immunostaining images of L3 SGs of the indicated genotypes. Centrosomes in yellow, mCherry tagged proteins in red and DNA in blue. The white squares mark the regions shown at higher magnification views in the middle panel. On the right, the centrosome channel is shown in grey. Pink dashed lines surround SG cells and yellow arrows point at centrosomes. (C and F) Bar and dot plot graphs showing the percentage of SGs cells with centrosomes of the indicated genotypes. (D) On the left, representative images of SG cells in *MyoII^mu^*^t^ obtained by super resolution microscopy. Centrosomes (yellow) and DNA (blue). The white squares mark the regions shown at higher magnification views on the right panel. iMyoII=inactive MyoII; aMyoII= active MyoII; Statistical significance is shown and determined by ordinary one-way ANOVA test. Bars show the mean ± SD. For (C) and (F) three and four experiments respectively were quantified with at least 10 SGs analyzed per condition.

While characterizing MyoII^mut^ SGs, we observed a clear reduction in both cell size and nuclear size (Supplementary Figure 3A), suggesting defects in ploidy levels. This was further supported by Fluorescent in situ Hybridization (FISH) analysis, which confirmed reduced ploidy (Supplementary Figure 3B–D). Given these broader cellular defects, we next asked whether we can distinguish MyoII requirements. To do so, and to study MyoII function, without the confounding effects of overexpression, we generated transgenic fly lines expressing MyoII under the control of its native promoter.

Expression of WT MyoII and inactive MyoII (iMyoII), a MyoII mutant form that cannot be phosphorylated due to alanine substitutions at the phosphorylation site (A21A22 to replace S21T22) (*32*), was analyzed in the *MyoII^mut^* background. Cell and nuclear size defects were rescued to values comparable to WT SGs in both WT MyoII and iMyoII (Supplementary Figure 3A). However, while WT MyoII SGs displayed an impressive decrease in the number of cells containing centrosomes, SGs expressing iMyoII, maintained centrosomes (Figures 2B-C). Both WT MyoII and iMyoII SGs presented centrosomes in the lumen, even though at a decreased frequency in iMyoII compared to WT SGs (Supplementary Figure 2B-C). These data show that MyoII activity is required to promote centrosome expulsion from SG cells. Moreover, they show that failure to remove centrosomes from SG cells leads to centrosome amplification.

### Centrosome release into the SG lumen does not require loss of Polo kinase activity

In *Drosophila* female oocytes, centrosome elimination occurs in a stepwise manner, beginning with the removal of the pericentriolar material (PCM) that surrounds the centrioles. This initial step depends on the de-localization of Polo (Polo-like kinase 1) away from the centrosome (13). To test whether a similar mechanism operates upstream of centrosome release into the SG lumen, we expressed Polo fused to the centriole-targeting domain PACT (Polo-PACT), which blocks PCM removal (13). Interestingly, in SG cells expressing Polo-PACT, an interference with centrosome expulsion was not seen (Supplementary Figure 3E–G). These results suggest that, unlike in oocytes, Polo loss and PCM shedding are not required for centrosome release in SG cells.

### Increasing MyoII activity can overcome the presence of extra centrosomes in SG cells

To test if increasing MyoII activity can overcome defects in centrosome expulsion we took advantage of Sak/PLK4 over-expression (Plk4OE), which leads to centrosome amplification in many different cells and tissues, leading to tumorigenesis (*27*, *35–39*). Analysis of Plk4OE SGs at L3 stages, showed cells containing centrosomes (Figures 2E-F and Supplementary Figures 1D-E), contrary to WT cells. Further, centrosomes were also detected in the lumen (Supplementary Figure 2E-F), suggesting that a certain number of centrosomes was nevertheless released into the lumen in Plk4OE SGs, indicating that the mechanism of elimination is likely present in Plk4OE SGs.

We next investigated whether increasing MyoII contributed to decrease the number of centrosomes in Plk4OE SG cells. Using the same strategy as described previously for WT MyoII and iMyoII, we expressed under its endogenous promoter, a constitutively active MyoII (aMyoII) transgene, where the S21T22 residues were replaced by glutamic acid residues (E21E22) mimicking phosphorylation (*32*). Importantly, this fully rescued the defects in centrosome elimination of Plk4OE SGs, while WT MyoII or iMyoII had no effect (Figures 2E-F). We concluded that MyoII activity is essential to promote centrosome expulsion in SG cells. A possible interpretation of these findings is that centrosome release towards the lumen is limited by the levels of active MyoII.

### Luminal centrosomes are targeted for proteasome degradation

We next investigated the fate of luminal centrosomes in WT SGs, using two centrosome markers to detect co-localization and therefore identify bona fide centrosomes (*35*, *40*, *41*). At L3 stages, a variety of centrosome sizes and structures could be detected in the WT SG lumen (Supplementary Figures 4A-C). These included centrioles, like the ones detected in other tissues (*34*) and centrosome filaments, where beads like structures detected by centrosome markers appeared clustered in a filament-like pattern. Finally, extremely small structures were also seen. We called these centrosome dust, in light of their small size. At later developmental stages however, when the lumen becomes enlarged, no centrosome structures could be detected, suggesting the disappearance of centrosomes as larvae transition towards the pupal stage (Supplementary Figure 4D-E).

These results suggest an ongoing process of centrosome degradation. Recently, it has been shown that in *Naegleria gruberi,* centrosomes undergo ubiquitination, which promotes proteasome-mediated degradation (*42*, *43*). To ascertain whether ubiquitination plays a role in promoting the disappearance of luminal centrosomes, we labelled WT SGs with antibodies that recognize ubiquitin moieties (P4D1) and K48 polyubiquitin (K48-Ub). In WT SGs, both markers co-localized with luminal centrosomes (Supplementary Figure 4F-H), suggesting indeed that these represent proteasome substrates (*44–46*). In *MyoII^mut^* SGs however, only P4D1 co-localized with centrosomes in cells, whereas no K48-Ub signal could be identified in the lumen (Supplementary Figure 4F-H).

Proteasome-mediated degradation within the lumen necessitates the localization of the proteasome to this compartment. Several studies have described the presence of the proteasome in extracellular space such as blood, extracellular alveolar space or in the cerebrospinal fluid (*47–50*). Nevertheless, the presence of the proteasome in the lumen of SGs, has never been reported. We therefore used a *Drosophila* proteasome-activity reporter line, where the proteasome substrate-CL1-is fused to GFP (*51*), so that in the presence of the proteasome, GFP fluorescence is not detected. At later developmental stages, GFP positive signals could be detected in a few SG cells, but not in the lumen (Supplementary Figure 4I), suggesting proteasome activity in the lumen.

To functionally test the possible role of the proteasome in L3 WT SGs, we used the proteasome inhibitor MG132 (*52–54*) for three hours, to avoid toxic effects. Importantly, this short incubation time was sufficient to completely alter the morphology of luminal centrosomes, which now could be seen forming large aggregates and filling the lumen entirely (Supplementary Figure 4J-K). Altogether, these results show that ubiquitinylated centrosomes are present in the SG lumen at late larval stages, which triggers their degradation in a proteasome dependent manner.

### Secretory autophagy coupled with autolysosome exocytosis promotes centrosome expulsion from SG cells

With the aim of identifying the mechanism responsible for centrosome expulsion from the cell into the lumen, we performed a candidate screen, where mutations or RNA interference (RNAi) of candidate genes were tested. We reasoned that if a given process involved in centrosome luminal release was inhibited, centrosomes should be detected within SG cells at later larval stages, as observed in *MyoII^mut^* SG*s* and contrary to WT *SGs*. We tested components known to be involved in the biogenesis of MVBs and therefore altering their secretory properties, such as the endosomal sorting complexes required for transport (ESCRT) proteins (*55–57*) and common secretory pathways (*58*).

Depletion of different ESCRT members or members of the secretory pathway (see methods) did not show any relevant difference in terms of centrosome behavior: L3 SG cells did not contain centrosomes, and as in WT, they could be detected in the lumen (Supplementary Figure 5A-C). It is worth noticing that SGs depleted for these components presented abnormal organization of the polytene chromosome in the nucleus, attesting for an effect of the RNAi.

We next tested autophagy using an amorphic mutation in the *atg5* gene (*Atg5^mut^*), which encodes a member of the autophagy complex, responsible for phagophore formation (*59*). Striking, similarly to *MyoII^mut^* or Plk4OE SGs, *Atg5^mut^* SGs cells still contained multiple centrosomes (Figure 3A-C), suggesting a requirement for this pathway in centrosome elimination in SG cells. Further, this supports the findings describe above showing that if centrosomes are not eliminated from SG cells, they keep duplicating.

**Figure 3.**
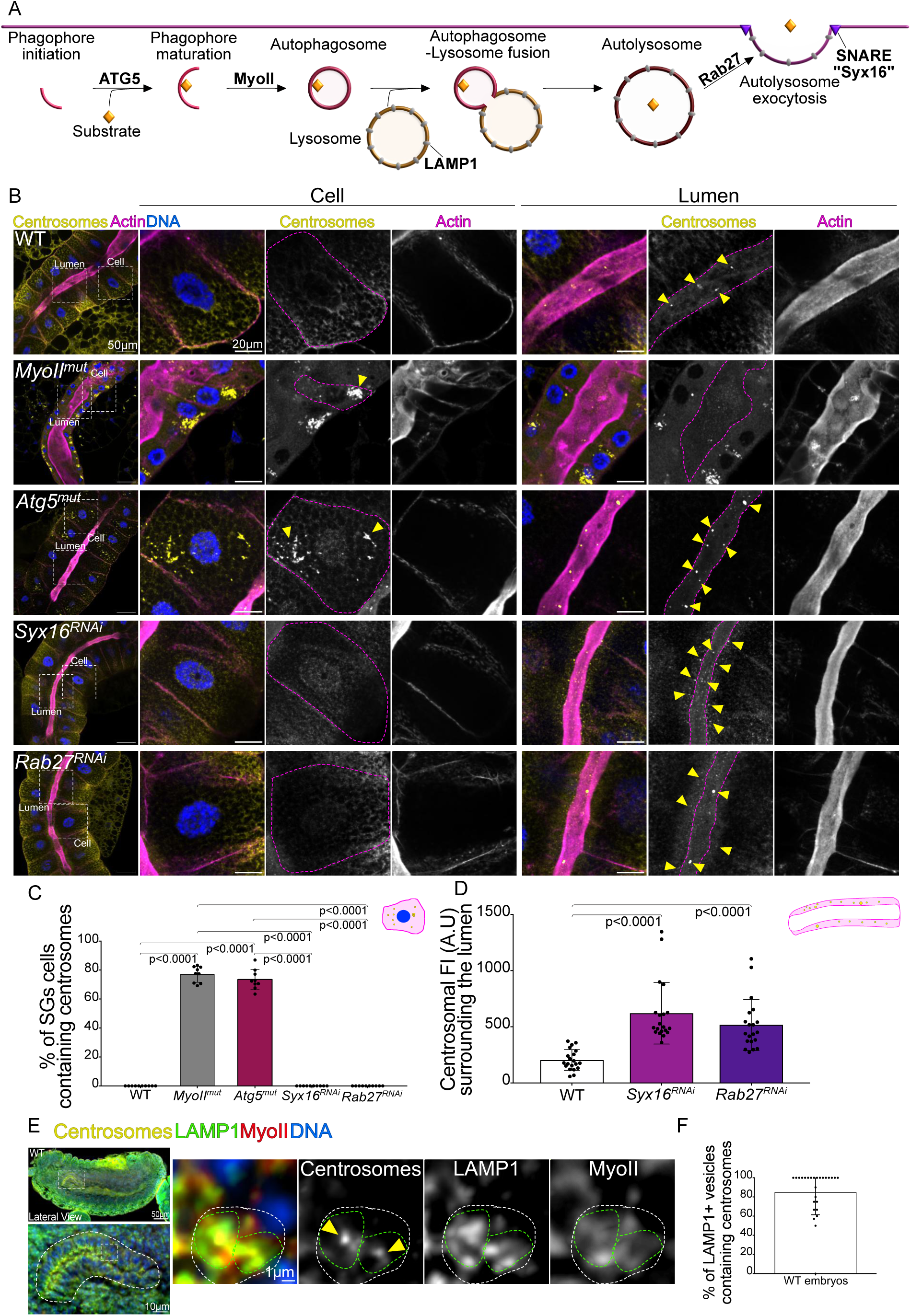
Secretory Autophagy and Lysosomal Exocytosis Mediate Centrosome Delivery into the Lumen. (A) Schematic diagram of the main steps involved in autophagosome formation and lysosomal exocytosis. (B) On the left, representative low magnification immunostaining images of L3 SGs of the indicated genotypes. Centrosomes in yellow, actin in magenta and DNA in blue. The white squares mark the two SG regions that are shown at higher magnifications on the right as merged panels or as separate panels showing centrosomes and actin as indicated.These regions correspond to a cell (middle panels) and the lumen (right panels). The dashed pink lines surround the cell, or the lumen and the yellow arrows point at centrosomes. (C-D) Bar and dot plot graphs showing the percentage of SGs cells with centrosomes (C) and the centrosomal fluorescence intensity (FI) in A.U. taken in the apical membrane and luminal regions as illustrated with the drawing on the top right corner. (E) On the left, representative immunostaining image of one embryo (top) and SG (bottom) showing centrosomes in yellow, LAMP1 in green, MyoII in red and DNA in blue. The white dashed rectangle and square show the SG and the SG region, shown at higher magnification on the bottom and on the right respectively. The white dashed square surrounds the LAMP1 vesicles and the green dashed lines surround the centrosomes. (F) Bar and dot plot graph showing the percentage of LAMP1+ vesicles containing centrosomes in WT embryos of stage 13-14. Statistical significance is shown and it was determined by ordinary one-way ANOVA test (C-D) test. Bars show the mean ± SD. For (C), three experiments and for (D-F) two experiments were quantified with 10 SGs in (C-D) and 28 embryos in (F).

Selective autophagy has been shown to regulate centrosome number in cells that normally contain centrosomes (*43*, *60*, *61*), by degrading specific centrosomal proteins such as Cep63 (*62*). Because in SGs, centrosomes are not degraded inside the cell but rather in the lumen, whereas in *Atg5^mut^* SGs centrosomes are retained within the cell, we hypothesize that autophagy here may function not as a degradative process but rather as a secretory one. Indeed, autophagy can support secretory processes, such as autolysosomal exocytosis (*63–65*). This process occurs through fusion of autophagosomes with lysosomes and subsequently with the plasma membrane, so that cellular content is released into the extra cellular space (Figure 3A).

To test the possible involvement of autolysosome exocytosis, we depleted Rab27 and Syntaxin (Syn)16. Rab27 is a small GTPase promoting lysosome docking at the plasma membrane, facilitating lysosomal exocytosis (Figure 3A) (*66*, *67*). Syx16, is a Soluble N-ethylmaleimide-sensitive factor attachment protein receptors (SNARE) protein involved in autolysosome formation (Figure 3A) (*68*). Remarkably, depletion of Rab27 or Syx16 showed centrosomes positioned closely to the apical membrane (Figure 3B-D), suggesting failure in the fusion process.

One common way of detecting autolysosome exocytosis is by monitoring the localization of lysosomal associated membrane protein 1 (LAMP1), which is a key protein involved in lysosome docking at the cell membrane (Figure 3A) (*69*, *70*). Using LAMP1 antibodies, we characterized WT SGs, at embryonic stages 15-17, which corresponds to the stages where centrosomes are released into the lumen. Importantly, we found a high enrichment of LAMP1 vesicles at the apical membrane of SGs (Figure 3E). Strikingly, these vesicles also contained MyoII and centrosomes (Figure 3E-F). These results support the findings that MyoII activity, combined with secretory autophagy and autolysosome exocytosis cooperate to expulse centrosomes from SG cells towards the lumen.

### Centrosome elimination is required for proper mitochondria organization and respiratory activity

A key unanswered question in the field of polyploidy is the underlying reason for centrosome elimination in certain polyploid cells. An obvious possibility is that the maintenance of centrosomes, that continue duplicating, may impact SG development or ploidy levels. *MyoII^mut^* SG cells display reduced size and ploidy, when compared to WT SG cells (Supplementary Figure 3A-D). However, Plk4OE SG cells contained centrosomes but normal ploidy levels and cell and nuclear size, when compared to WT (Supplementary Figure 3A-D), showing that lack of centrosome expulsion does not impair SG cell growth or increase in ploidy.

We next hypothesized that the presence of centrosomes may hinder MT nucleation capacity from the nuclear envelope, typical of SG cells (*71*). Indeed, in mammalian or *Drosophila* muscle cells or in the *Drosophila* oocyte (*72–74*), MTs are nucleated from the nuclear envelope in the absence of centrosomes and form a nuclear ring. Whether centrosome presence and nucleation from the nuclear envelope are mutually exclusive is unknown. In WT larval SGs, a highly organized MT ring could be readily identified. In sharp contrast, *MyoII^mut^ SGs* lacked MT rings and displayed a very disorganized MT network (Supplementary Figure 6A-B). However, analysis of Plk4OE SGs showed an intact and perfectly arranged MT ring in cells that contain centrosomes (Supplementary Figure 6A-B). These results indicate that in polyploid cells, MTs can be nucleated by the nuclear envelope even when centrosomes are present. This suggests that the primary purpose of centrosome elimination in polyploid cells is not to enable MT nucleation from alternative MTOCs.

Recently, it has been shown in human cells that the presence of extra centrosomes-centrosome amplification-induces defects in mitochondria organization and activity (*75*, *76*). Further, other studies have shown that in specialized cells like neurons or fibroblasts, MT motors regulate mitochondria motility and activity and defects in MT-dependent mitochondria transport can impair the function of this organelle (*77*, *78*). Using electron microscopy (EM) approaches we found that mitochondria appeared smaller in *MyoII^mut^* SG larval cells when compared to WT cells (Figure 4A-B). Further, using parameters such as circularity and solidity to infer on the general morphology aspect and branching respectively, showed elevated indices in MyoII^mut^. Motivated by these findings, we next labelled the mitochondrial network in WT, *MyoII^mut^*and Plk4OE SGs using antibodies against the mitochondrial ATP synthase F1 Subunit Alpha (ATP5a). Interestingly, in WT SG cells, mitochondria appeared filamentous and formed a long and branched network (Figure 4C). In both *MyoII^mut^*and Plk4OE SG cells however, mitochondria appeared smaller and fragmented with reduced surface area (Figure 4D). Analysis of the circularity and solidity parameters showed higher values in both *MyoII^mut^* and Plk4OE SG cells, when compared to WT SGs (Figure 4E-F), confirming a higher degree of branching in WT cells. These results suggested that maintaining centrosomes in SGs impact mitochondria structure and network organization.

**Figure 4.**
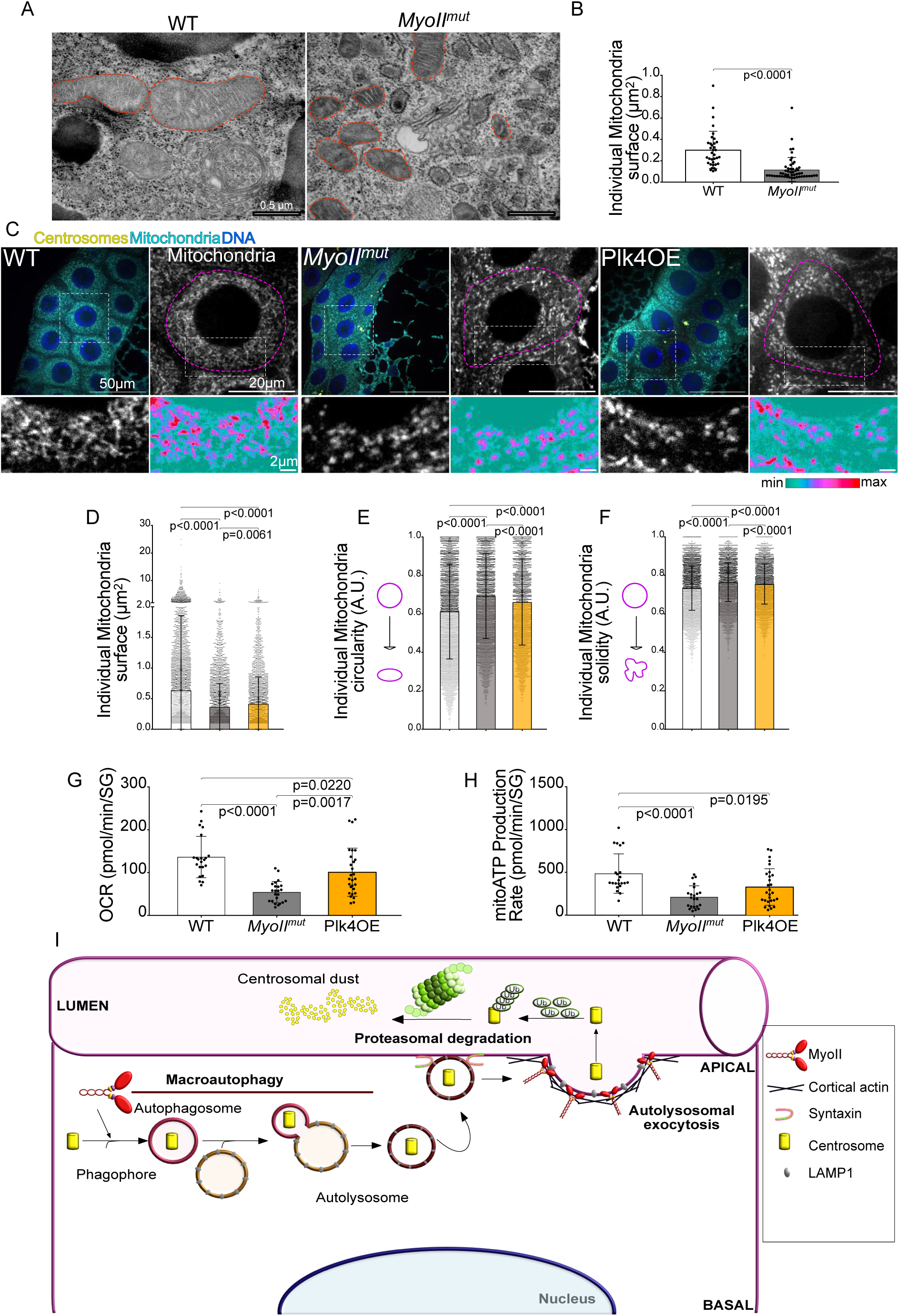
Maintenance of centrosomes in SGs cells impacts the mitochondrial network and activity. (A) Representative EM images of the indicated genotypes. Orange dashed lines surround mitochondria. (B) Bar and dot plot graphs showing individual mitochondria surface from EM data of the indicated genotypes. (C) Representative immunofluorescence images of L3 SGs of the indicated genotypes showing centrosomes in yellow, mitochondria in cyan and DNA in blue on the left and mitochondria on the right in grey and in higher magnifications in grey and on the Ice LUT bellow. The white dashed squares show the magnification regions. The pink dashed lines surround SG cells. (D-F) Bar and dot plot graphs showing individual mitochondria surface (D), circularity (E) and solidity (F) of the indicated genotypes. (G-H) Bar and dot plot graphs showing oxygen consumption rates (OCR) (G) and ATP production rates from mitochondria (H) in SG cells of the indicated genotypes. (I) Model representing the main findings of this study. During embryonic stages, MyoII functions together with the autophagy machinery and lysosomal exocytosis to transport centrosomes into the lumen of SGs. Centrosomes are then ubiquitinated and degraded by the proteasome machinery at later developmental stages. Statistical significance is shown and determined by Mann-Whitney test (B) and ordinary one-way ANOVA tests (D-F and G-H). Bars show the mean ± SD. For (B) a minimum of 35 mitochondria were analyzed from two independent experiments. For (D-F), two experiments with a minimum of 10 SGs were analyzed and for (G-H), a minimum of 22 SGs were analyzed per condition.

Mitochondria are key organelles that produce energy through ATP synthesis (*79–82*). Cells are thought to increase their ploidy to enable larger cell size, which may be linked to higher metabolic activity. We explore this possibility by measuring oxygen consumption rates (OCR) and mitochondrial ATP production rates using the Seahorse XF Analyzer. Strikingly, an important decrease in both OCR and mitochondrial ATP production rates was detected in *MyoII^mut^* and Plk4OE SG cells, compared to the levels seen in WT SG cells (Figure 4G-H). Altogether, our findings uncovered a novel role for centrosome elimination in ensuring proper mitochondria organization to maintain metabolic and respiratory activity in polyploid SG cells.

## Discussion

Our findings highlight the essential role of centrosome elimination in physiologically polyploid cells. They support a model (Figure 4I) in which centrosome removal depends on MyoII activity, which facilitates the transport of centrosomes from embryonic SG cells into the lumen. MyoII is known to function in autophagy, particularly in autophagosome formation (*83*), and our data show that inhibition of either MyoII or autophagy leads to the retention of centrosomes within the cell. Based on our results, we propose that autophagosomes containing centrosomes fuse with lysosomes to form autolysosomes, which are subsequently directed towards the plasma membrane, where they can undergo membrane fusion in a Syx16- and Rab27-dependent manner. This enables the release of centrosomes into the lumen, where they are eventually degraded by the proteasome during later developmental stages.

The strict requirement for complete centrosome elimination in these and other polyploid cells has remained unclear. Our findings suggest that centrosome retention disrupts mitochondrial network integrity, compromising the cell’s ability to generate ATP at necessary levels. It remains unknown whether luminal centrosomes serve any biological function before their degradation.

Interestingly, recent studies have reported that human prostate and ovarian tumors often exhibit centrosome loss (41, 84). Although it remains unclear whether this phenomenon is linked to changes in mitochondrial respiratory activity, our results, together with these findings, highlight the importance of investigating centrosome loss to understand the potential cellular advantages it may confer.

How ectopic presence of centrosomes affect mitochondria function remains to be understood. One possibility is that amplified centrosomes alter the polarity or properties of the MT cytoskeleton, which may impact fusion explaining the decrease in mitochondria size observed. Alternatively, the presence of extra centrosomes may influence other aspects of mitochondria biogenesis and function that remain to be understood. Importantly, this work uncovers a physiological requirement for centrosome elimination in cells undergoing extensive ploidy and cell size increase, particularly in support of large-scale metabolite production.

## Acknowledgments

The authors acknowledge the Cell and Tissue Imaging (PICT-IBiSA) from Institut Curie, member of the national infrastructure France-BioImaging (https://ror.org/01y7vt929) supported by the French National Research Agency (ANR-24-INBS-0005 FBI BIOGEN) and V. Fraisier, C. Guedj. M. David, M. Cortes for their continuous support on image acquisition and analysis. We Thank A. Martin (MIT) for providing the Sqh (MyoII promoter). The authors thank B. Goud, F. Perez, Y. Bellaiche, J.L. Maitre (I. Curie), W. Marshall (UCSF), G. Montagnac (IGR) and M. Thery (IPGG) for helpful discussions. We are grateful to A.C. Gavin Perrin (UNIGEN) for bringing the lysosomal exocytosis pathway into our attention and for A. Echard for guidance throughout the project. We thank J. Mathieu, J. Schulman and U.B. Pandey for *Drosophila* stocks. We thank V. Marthiens and A. Echard for comments on the manuscript. This work has been supported by the European Research Council ERC CoG (ChromoNumber-LS3, ERC-2016-COG for R.B.), the government through the Agence Nationale de la Recherche from France 2030 (ANR-23-CHBS-0012 for R.B), the Institut Curie and the CNRS. R.S salary was supported by the ERC-2016-COG and an FRM grant (FDT202404018667).

## Author’s contribution

R.S. and R.B. conceived the project and wrote the manuscript. R.S. did most of the experiments and data analysis presented here with some help from A. S. L.J and S.G P.B. and G.R. contributed with the EM experiments and C.C with the seahorse measurements. R.B. obtained the funding used to develop this project. All authors read and commented on the manuscript.

## Materials and Methods

### Fly culture

For all experiments performed on larvae, flies were raised in plastic vials containing homemade standard *Drosophila* rich culture medium (0.75% agar, 3.5% organic wheat flour, 5% yeast, 5.5% sugar, 2.5% nipagin, 1% penicillin-streptomycin (Gibco #15140), and 0.4% propanic acid). Fly stocks and experimental crosses were maintained at 22°C. Stocks were maintained using balancer inverted chromosomes to prevent recombination. In all experiments, larvae were staged to obtain comparable stages of development. Egg collection was performed at 22°C for 24 hours (h). Mid-Late third instar larvae (L3) salivary glands (SGs) were dissected after 7 days of development. Pre-pupae larvae were dissected after 9 days (by choosing immobilized larvae on the wall of the tube).

### Fly stocks

BL = Bloomington Drosophila Stock Center (Indiana University, IN, USA).

| Stock Name | Chromosome | Source | Reference |
| --- | --- | --- | --- |
| wild-type (WT) - w1118 |  | BL3605 |  |
| **Driver** |  |  |  |
| Fkh-Gal4 | II | BL78061 | (1) |
| **Fluorescent proteins** |  |  |  |
| Fkh-GFP | II | BL43951 | (2) |
| Sas-4 – GFP | II | Raff lab | (3) |
| MyoII-mCherry (S20-T21) | III | Basto lab | This work |
| iMyoII-mCherry (A20A21) | III | Basto lab | This work |
| aMyoII-mCherry (E20E21) | III | Basto lab | This work |
| UAS-CL1-GFP | II | Pandey lab | (4) |
| UAS-Polo-PACT-GFP | II | Bettencourt-Dias lab | (5) |
| **Mutants** |  |  |  |
| *sqh*<sup>1</sup> (*MyoII*<sup>mut</sup>) | X | Karess Lab | (6) |
| *atg5*<sup>5cc5</sup> (*atg5*<sup>mut</sup>) | X | BL91419 | (7) |
| *alix*<sup>1</sup> (*alix*<sup>mut</sup>) | III | BL85211 - J.Mathieu lab | (8) |
| Plk4OE | III | Raff lab | (9) |
| **RNAi Lines** |  |  |  |
| UAS-Syx16-RNAi | III | BL25884 | (10) |
| UAS-Rab27-RNAi | III | BL35774 | (11) |
| UAS-Chmp2b-RNAi | II | BL38375 | (12) |
| UAS-Shrub-RNAi, UAS-Chmp2b-RNAi | III | Huynh lab | (8) |
| UAS-Rab6-RNAi | III | BL35744 | (13) |
| UAS-Rab7-RNAi | III | BL27051 | (10, 13) |

### Immunofluorescence of *Drosophila* SGs whole mount preparations

L3 SGs were dissected in fresh PBS and fixed for 40 minutes (min) at room temperature (RT) in 4% paraformaldehyde (EMS #15710) in PBS. Fixed tissues were then washed 3×15min in PBST 0.3% (PBS + 0,3% Triton X-100, Euromedex #2000-C) and incubated for 1h with blocking solution (PBST + 5% normal goat serum (NGS), Gibco #10098792). After washing 3×15min in PBST, tissues were incubated in primary antibodies at the appropriate dilution in the blocking solution, overnight at 4°C in a humid chamber. After washes in PBST 3×15min, tissues were incubated in secondary antibodies diluted in the blocking solution, protected from light in a humid chamber, for 4h at 25°C. Tissues were then washed with PBST 3×15min, rinsed in PBS and incubated with DAPI (4′,6-diamidino-2-phenylindole; 1:1000, Thermo Fisher Scientific #D1306) for 30 min, protected from light. SGs were then rinsed in PBS and mounted in clearing solution (Rapiclear 1.49 Sunjin Lab #RC149002).

### Clearing experiment on L3 SGs for microtubule staining

For labelling the microtubule (MT) cytoskeleton, we used a clearing protocol that permeabilizes and improves antibody penetration generating tissues with lower fluorescence background levels. L3 SGs were fixed with 4% paraformaldehyde for 3h on a shaker at RT, then washed with PBS 4×30min. Tissues were transferred into a PBST 2% solution (PBS and 2% Triton X-100) on a shaker for 2h at RT for permeabilization and then washed again with PBS for 4×30min. The solution was changed to a freshly prepared blocking buffer with 10% NGS, 1% Triton X-100, 2.5% DMSO (Sigma #276855) and 0.2% sodium azide (Sigma #71289) in PBS-on a shaker for 24h at 4°C in a humid chamber. In the following day, primary antibodies were added to the antibody buffer-1% NGS, 0.2% Triton X-100, 2.5% DMSO and 0.2% sodium azide in PBS. Samples were placed on a shaker for 24h at 4°C in a humid chamber. Samples were washed with a freshly autoclaved washing buffer 3% NaCl (Honeywell Research Chemicals #31434) and 0.2% Triton X-100 in PBS-3×1h at RT and then incubated with secondary antibodies also diluted in antibody buffer. Samples were placed on a shaker for 24h at 4°C in a humid chamber. They were washed again with PBS 4×30min at RT. Tissues were then rinsed in PBS and incubated with DAPI (1:1000) at RT for 1h, rinsed again with PBS and incubated in the clearing solution for 1h before being mounted in the clearing solution.

### Embryo culture and collection

Embryos were collected following ON egg laying on agar apple juice plates. Embryos were fixed either using formaldehyde or methanol. The use of Formaldehyde fixative allows phalloidin labeling. After collection with a small amount of detergent solution (PBST 0.05%), embryos were transferred to a clean Eppendorf before being dechorionated in 50% freshly diluted bleach for 3min, then washed extensively with water and fixed in 2:1 mixture of Heptane and 4% methanol-free formaldehyde (EMS #15686) for 25min and devitellinized in a 50% mix of 90% ethanol (Avantor Sciences #20821.310) and heptane (Dutscher #446951) with vigorous shaking for 30 seconds (sec). Embryos were then rinsed twice with PBT 0.3% and incubated for 1h at 4°C with rotation in blocking solution-0.5% BSA (Euromedex #04-100-812-C) + 0.3% Triton X-100 in PBS. They were then labelled with phalloidin and primary antibodies in the blocking solution, ON at 4°C with rotation. The next day, embryos were washed with blocking solution 3×30min at RT and secondary antibodies were added to the blocking solution for 3h at RT, protected from light with rotation. Embryos were washed with the blocking solution 3×20min at RT, rinsed twice with PBS, incubated in DAPI (1:1000) for 20min at RT, protected from the light, and then mounted in a homemade mounting medium HMMM (1.25% n-propyl gallate (Sigma #P3130), 75% glycerol (bidistilled, 99.5%, VWR #24388-295), 23.75% H2O).

For methanol Fixation (used for Lamp1 staining), embryos were collected as before. Embryos were transferred into an Eppendorf with 1:1 mixture of heptane and methanol-EGTA solution (heptane, 97% methanol (Honeywell #603-001-00-X) and 3% 0.5M EGTA (Euromedex #1310-B), shaked for 30sec and then the solution was replaced with methanol. Embryos were stored at 4°C. For labelling, embryos were rehydrated with PBT 0.5% on a rotator 2×15min and then incubated in a blocking solution (5% BSA in PBS) 2×30min at RT on a rotator and incubated with primary antibodies diluted in the following blocking solution (5% BSA in PBST 0.05%), ON on a rotator at 4°C. The next day, embryos were washed with PBST 0.05% 3×20min at RT and then incubated with secondary antibodies diluted in the same blocking solution for 2h with rotation at RT protected from light. Embryos were then washed for 20min with PBST 0.05% at RT and incubated in DAPI (1:1000) for 20min at RT, protected from light and then mounted in HMMM.

### DNA Fluorescent In Situ Hybridization (FISH)

After fixation with 4% paraformaldehyde for 40min and washed with PBST 0.3% 3×15min, SGs were rinsed 3x in PBS, washed for 5min in 2xSSCT (2X Saline Sodium Citrate (Euromedex #EU0300-A) 0,1% Tween-20 (Sigma Aldrich #P1379), diluted in water, then for another 5min in 2xSSCT/50% formamide (Sigma Aldrich #47671) and transferred in pre-warmed 2xSSCT/50% formamide and pre-hybridized for 3min at 92°C. In the meantime, DNA probes diluted in the Hybridization Buffer (20%dextransulfate (Sigma Aldrich #D8906), 2xSSCT, 50%formamide, 0,5mg/ml salmon DNA sperm (Sigma Aldrich #D1626) were denatured at 92°C. After supernatant removal, SGs were incubated with the probe solution and hybridized for 5min at 92°C and 5h at 37°C. Samples were rinsed at RT, washed for 10min at 60°C in a 60°C pre-warmed 2xSSCT solution and for another 5min at RT in 2xSSCT. Finally, after rinsing in PBS, SGs were incubated in DAPI (1:1000) for 20min at RT and mounted in HMMM. DNA probes (Sigma-Aldrich) used in this study were designed to recognize sequences localized on chromosome II and III, coupled with Cy3 fluorescent dye (40ng/µl) and Cy5 fluorescent dye (80ng/µl), respectively as described in (Illuminati paper).

### Primary and secondary antibodies used in this study

| Antibody | Dilution | Source | Identifier |
| --- | --- | --- | --- |
| **Primary antibodies** |  |  |  |
| Guinea pig anti- Sas-4 | 1/500 | Raff J.W. Lab | (14) |
| Rabbit anti- Plp | 1/1000 | Raff J.W. Lab | (15) |
| Guinea pig anti- Cnn | 1/500 | Raff J.W. Lab | (16) |
| Mouse monoclonal anti-γTub | 1/500 | Sigma-Aldrich | T5326 |
| Rat monoclonal anti-RFP | 1/500 | Chromotec | 5F8 |
| Chicken polyclonal anti-GFP | 1/1000 | Abcam | ab13970 |
| Rabbit polyclonal anti-Lamp1 | 1/250 | Abcam | ab30687 |
| Mouse monoclonal anti-ATP5A (15H4C4) | 1/100 | Abcam | ab14748 |
| Rabbit polyclonal anti-aPKC (C-20) | 1/500 | Santa-Cruz | sc-216 |
| Mouse monoclonal anti-αTubulin | 1/500 | Sigma | T9026 |
| Rabbit Ub-K48 Recombinant Rabbit Monoclonal Antibody (5Y8J0) | 1/200 | Invitrogen | MA5-35382 |
| Mouse Ubiquitin monoclonal antibody (P4D1) | 1/500 | Enzo Life Sciences | BML-PW0930-0100 |
| Rabbit Anti-Myosin Light Chain Antibody, phospho-specific (Ser19) | 1/200 | EMB millipore | ab3381 |
| Alexa Fluor™ Plus 555 Phalloidin | 1/250 | Thermo Scientific | A30106 |
| Alexa Fluor™ Plus 647 Phalloidin | 1/250 | Thermo Scientific | A30107 |
| **Secondary antibodies** |  |  |  |
| Goat anti-Rat IgG (H+L) Cross-Adsorbed Secondary Antibody, Alexa Fluor 546 | 1/250 | Thermo Fisher Scientific | A11081 |
| Goat anti-Mouse IgG (H+L) Highly Cross-Adsorbed Secondary Antibody, Alexa Fluor 546 | 1/250 | Thermo Fisher Scientific | A11030 |
| Goat anti-Guinea Pig IgG (H+L)<br>Highly Cross-Adsorbed Secondary<br>Antibody, Alexa Fluor 488 | 1/250 | Thermo Fisher<br>Scientific | A11073 |
| Goat anti-Rabbit IgG (H+L) Highly<br>Cross-Adsorbed Secondary<br>Antibody, Alexa Fluor 647 | 1/250 | Thermo Fisher<br>Scientific | A21245 |
| Goat anti-Chicken IgY (H+L)<br>Secondary Antibody, Alexa Fluor 488 | 1/250 | Thermo Fisher<br>Scientific | A11039 |

### Live imaging of *Drosophila* embryos

A glue solution containing double side sticking tape diluted in heptane was prepared with rotation. The glue solution was then spread on a glass-bottom 35mm dish (MatTek Corporation #P35G-1.5-14-C) and once dried, embryos were placed on the plate and rolled on the glue to remove the chorion. Subsequently, embryos were positioned ventrally, facilitating visualization of SGs. Embryos were covered with oil 10S Voltalef (VWR BDH Prolabo). Images were acquired with 100x/1.4 NA oil objective, at time intervals of 30 sec and 20 to 40 Z-stacks of 0.2µm.

### MG-132 treatment for proteasome activity inhibition

L3 SGs were dissected in Schneider’s *Drosophila* medium used for live imaging (described above) and incubated for 3h at 25°C in Schneider’s medium with MG-132 20 µg/ml (Selleck Chemicals #S2619, diluted in DMSO). Control SGs were incubated for 3h at 25°C in Schneider’s medium with DMSO 20 µg/ml. Tissues were washed 3x in PBS and were fixed and stained as described above.

### SeaHorse Assays

Real-time ATP production rate was measured on a SeaHorse XFPro Analyzer. The day before, the SeaHorse cartridge was hydrated using calibrant (Agilent #100840-000) and placed in an incubator at 25^°^C without CO_2_. The SeaHorse was also set at 25°C. The morning of the experiment, L3 SGs were dissected and placed directly in the bottom of a SeaHorse 96-well plate (Agilent #103792-100) containing SeaHorse XF DMEM (Agilent #103575-100) supplemented with 10mM glucose (Agilent #103577-100), 1mM pyruvate (Agilent #103578-100) and 2mM glutamine (Agilent #103579-100). After measuring basal metabolism, a mitochondrial stress test was performed by successive injection of oligomycin 15µM (Sigma-Aldrich #O4876-5MG, stock at 30mM in ethanol) then rotenone 50mM (Sigma-Aldrich #R8875-1G, stock at 50mM in chloroform) and antimycin A 5µM (Sigma-Aldrich #A8674-25MG, stock at 0.5mg/mL in DMSO) combined together for the second injection. For each step, oxygen consumption rate (OCR), and the flux of protons-the extracellular acidification rate (ECAR) was measured during twelve cycles to ensure that steady-state values were reached once the drug had fully penetrated the tissue.

### Conventional transmission electron microscopy (EM) experiment

L3 SGs were dissected in Schneider’s *Drosophila* Medium used for live imaging (described above) and embedded in low-melting agarose (2% w/v, Thermofisher Scientific #16520050). Tissues were fixed in 2.5% glutaraldehyde (EMS, Cat #16120), in Cacodylate buffer 100mM pH 7.2 (EMS, Cat #12300), ON at 4°C and then rinsed 3x/5min in 100mM cacodylate buffer pH 7.2 at RT. Tissues were next incubated in 1% OsO4 (EMS, Cat #19150), 1.5% potassium ferrocyanure (Sigma #702587) in 100mMcacodylate buffer pH 7.2, 45 min at 4°C. Rinsed again 3×5min in H2O at RT and dehydrated at RT successively 15 min in 40% ethanol, 15 min in 70% ethanol, 15 min in 90% ethanol, 15min in 95% ethanol, 3 x 15min in 100% ethanol, 10 min in acetone. Infiltrate with EMbed 812 (EMbed 812 Embedding Kit, EMS cat #14120) resin successively with EPON 30% for 30min, epon 50 % for 30min, epon 70% for 1h and epon overnight. Samples are then deposited in a flat embedding mold filled with pure epon and set to polymerize for 48h in an oven at 60°C. Then, ultrathin epon sections (70 nm) of SGs were prepared with an ultramicrotome (UCT, Leica Microsystems), and contrasted with uranyle acetate and lead citrate (Delta Microscopies, ref 11300) before observation at an 80 kV transmission electron microscope (Tecnai Spirit G2; Thermo Fisher Scientific) equipped with a 4k CCD camera (Quemesa, Soft Imaging System).

### Molecular Biology

Three constructs containing the *spaghetti squash (sqh)* gene-here referred to as MyoII-with a m-Cherry in tag in frame at the C-terminus were synthetized and mutagenized by Genescript (Rijswijk, Netherlands). WT MyoII (S21T22) was synthetized from 5’ UTR to the residue before the stop codon. The mutant versions were generated using standard procedures. MyoII AA (inactive MyoII) where alanines replaced S21 and T22 and MyoII EE (constitutively active MyoII), where glutamic acids replaced S21 and T22 sequences in this order: The constructs contained from 5’ to 3’ the following restriction enzymes: SpeI-MyoII - Linker - mCherry - BamHI -3’. After confirmation of the full length sequences by Sanger sequencing, 2µg of all constructs were digested and ligated to previously digested pBlueScript II SK(+)vector containing the *Drosophila* MyoII endogenous promoter sequence - a kind gift by A. Martin. Selection of positive clones was done following standard procedures including Sanger sequencing by Eurofins Genomics (Ebersberg, Germany). All the inserts were inserted in a P[acman] plasmid using AscI and NotI restriction enzymes (New England Biolabs). After verification, one clone was used to generate transgenic flies by PhiC31 integrase-mediated transgenesis on ChIII-insertion site-9750 by BestGene (CA, United States). Sequences will be available upon request.

### Image acquisition and analysis

Image analysis and quantifications were performed using Image J software V2.1.0/1.53c, and Imaris software v.9.8.2 (Bitplane, RRID:SCR_007370).

### L3 SGs Image acquisition and super-resolution

L3 SGs images were acquired using a Nikon Ti2-E fully motorized inverted microscope equipped with a 40x/1.3 NA and 100×/1.4 NA Plan-Apochromat oil immersion objectives and a Prime95B sCMOS camera (Teledyne Photometrics). Optical sectioning was performed via a Nano-Z500 piezoelectric stage (Mad City Labs). Excitation was provided by a laser combiner (Gataca Systems) comprising 405, 491, 561, and 642 nm laser diodes (150 mW each), coupled to a Yokogawa W1 spinning disk confocal head through a single-mode fiber. Multi-dimensional image acquisition was carried out in streaming mode using MetaMorph software (version 7.10.5, Molecular Devices). For super-resolution imaging, the Live SR module (Gataca Systems) was used in conjunction with the W1 spinning disk system. This module is based on structured illumination with the optical reassignment technique and online processing leading to a two-time resolution improvement. The method called multifocal SIM (MSIM) allows combining the doubling resolution together with the physical optical sectioning of confocal microscopy. Acquired images were processed and reconstructed using Gataca’s proprietary software integrated within the MetaMorph environment.

### Embryonic SGs Images acquisition

Images of embryos were acquired using a Nikon Ti2-E fully motorized inverted microscope equipped with the Nikon AXR line scanning confocal and 20x/0.8 NA and 60x/1.42 NA objectives. Optical sectioning was performed via a NanoScan SP Z 600um piezoelectric stage (Prior). Excitation was provided by Nikon laser combiner comprising 405, 488, 561, and 640 nm laser diodes (150 mW each). Multi-dimensional image acquisition was carried by resonant scanner with NIS software (version 6.10.01 (Build 2027) Patch 01, Nikon). Acquired images were processed by the “Denoise.ai” module of Nikon.

### Imaris

For the 3D reconstruction, images were converted in Imaris files using ImarisFileConverter 9.9.1 (Bitplane, RRID:SCR_007370). Then, images were imported into Imaris software v.9.8.2 (Bitplane, RRID:SCR_007370). 3D views were generated using Imaris software with surface module and saved as TIFF and mp4.

### Quantification of the frequency of events

Image analysis and quantifications were performed using Fiji software. Centrosomes in the embryonic SGs, were manually counted following the signal of Sas-4 and their localization was manually determined and confirmed by 3D reconstructions. Centrosomes were classified as being positioned in the cell, at the apical membrane (when in close proximity with the apical membrane) or in the lumen.

In L3 SGs, the number of cells with or without centrosomes was counted manually by identifying centrosomes using two centrosomal markers (Plp and Sas-4).

### Characterization of centrosomes in L3 SGs

To quantify the presence/absence or distribution of centrosomes in the SG lumen, a max projection was performed on all slices comprising the lumen. The luminal region was selected using the polygon selection tool. On both Sas-4 and Plp channels, centrosomes were identified, and their surface area was measured using the 3D object counter mode with adjusted threshold. For the centrosomes that were localized around or near the lumen in Rab27-RNAi and Syx16-RNAi conditions, a max projection was generated as before. Then, using the line tool, the area around the lumen was selected and the fluorescence mean intensity was measured for both centrosomal markers (Plp and Sas-4).

## Mitochondria characterization

### From immunostained data

SG cells were cropped using a macro adapted from Anne-Sophie Mace, a research engineer at the Institut PICT platform. The macro selected the center plane of the cell, and mitochondria were identified using the analyze particle module on Fiji. Parameters like area, circularity and solidity (two indices informing on object circularity and invaginations, respectively) were obtained with the measurement tool from ImageJ.

### From EM data

Mitochondria were identified in EM images based on their typical structure. Using the polygon selection tool and the measurement tool from ImageJ, parameters like area, circularity and solidity were obtained.

### Ubiquitin fluorescence signal quantification

Centrosomes were identified either in the cell (*MyoII^mut^*) or in the Lumen of WT SGs, with two centrosome markers (Plp and Sas-4). Each centrosome was selected using the polygon selection tool, and the mean intensity of both ubiquitin markers was measured.

### Statistical analysis

The statistical significance was calculated using GraphPad Prism (RRID:SCR_002798) version 7 for Mac, GraphPad Software, La Jolla California USA, www.graphpad.com. The statistical test used for each experiment is indicated in the figure legends. Each representative image originates from the analyzed dataset and are representative of the associated quantifications.

