## Supplementary Figures and Figure legends for "A novel mechanism for centrosome expulsion ensures metabolic activity in polyploid cells"

### Supplementary Figure 1

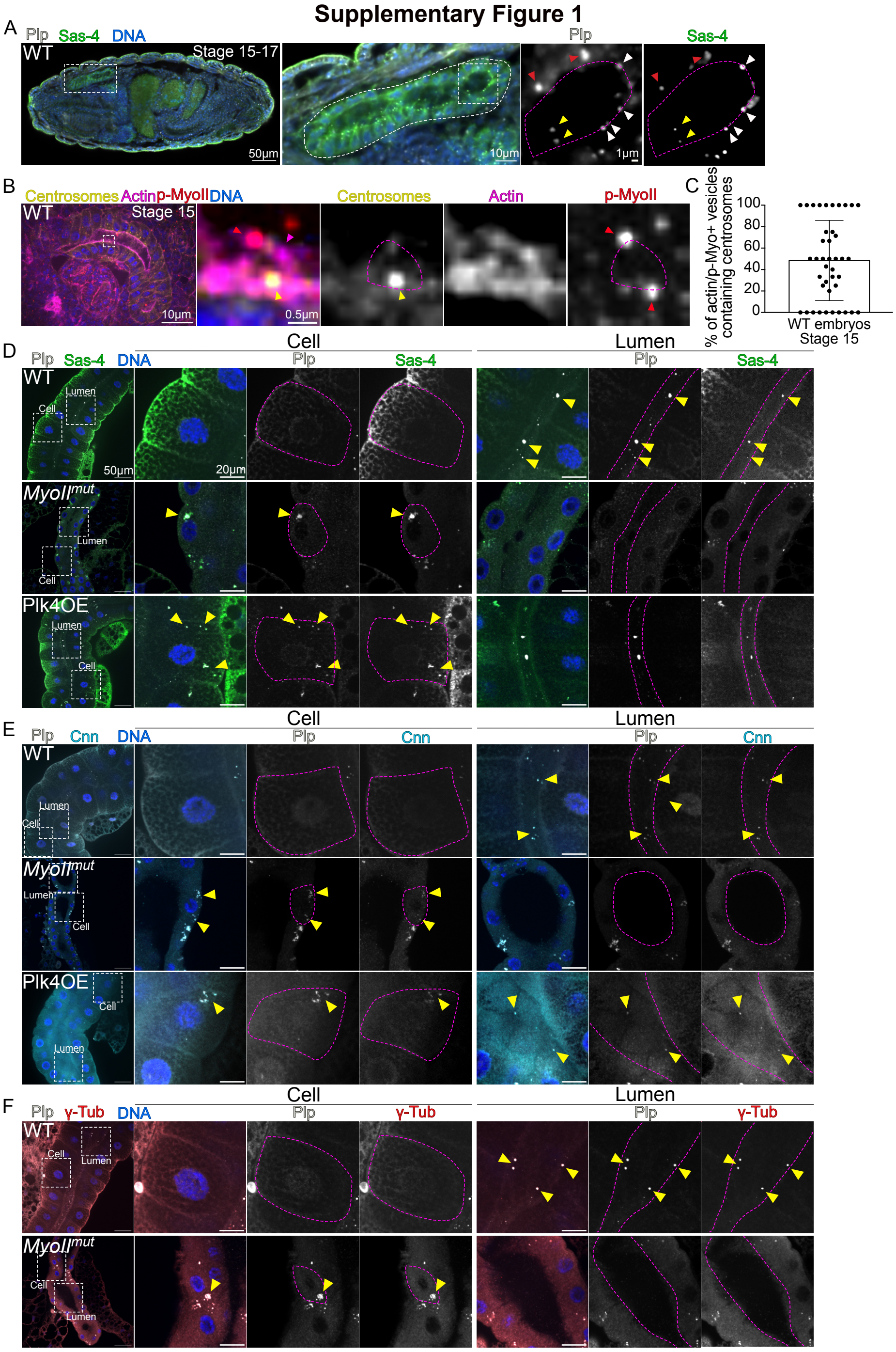

**Supplementary Figure 1- Validation of centrosome labeling with the use of two different centrosomal markers**

(A) Representative immunofluorescence images of WT embryos at stage 15-17 showing the co-localization of two centrosome markers- Plp in grey and Sas-4 in green. The white dashed square show the insets shown at higher magnifications on the right. The pink dashed lines surround SG cells. Yellow arrowheads point to centrosomes in the lumen, white arrowheads to centrosomes at the apical cell membrane and red arrowheads at centrosomes inside the cell. (B) Representative immunofluorescence images of embryonic SGs showing centrosomes in yellow, Actin in magenta and p-Myo in red. DNA in blue. The white dashed square shows the inset shown at higher magnifications on the right. The pink dashed line surrounds the actin vesicle. The yellow arrowhead point at a centrosome and the red arrowheads point to p-MyoII foci. (C) Bar and dot plot graph showing the percentage of actin/p-Myo+ vesicles containing centrosomes. (D) On the left, representative low magnification immunostaining images of L3 SGs of the indicated genotypes. Centrosomes are shown through the localization of two centrosome markers- Plp (grey) and Sas-4 (green), DNA in blue. The white squares mark the two SG regions that are shown at higher magnifications on the right as merged panels or as separate panels showing centrosome markers in grey. These regions correspond to a cell (middle panels) and to the lumen (right panels). The pink dashed lines surround SG cells or the lumen. Yellow arrowheads point at centrosomes. (E) On the left, illustrative low magnification immunostaining images of L3 SGs of the indicated genotypes. Centrosomes are shown through the localization of two centrosome markers- Plp (grey) and Cnn (Cyan) with DNA in blue. The white squares mark the two SG regions that are shown at higher magnifications on the right as merged panels or as separate panels showing centrosome markers in grey. These regions correspond to a cell (mid panels) and to the lumen (right panels). The pink dashed lines surround SG cells, or the lumen. Yellow arrowheads point at centrosomes. (F) On the left, illustrative low magnification immunostaining images of L3 SGs of the indicated genotypes. Centrosomes are shown through the localization of two centrosome markers- Plp (grey) and $\gamma$-tubulin ($\gamma$-tub) (red). DNA in blue. The white squares mark the two SG regions that are shown at higher magnifications on the right as merged panels or as separate panels showing centrosome markers in grey. These regions correspond to a cell (mid panels) and to the lumen (right panels). The pink dashed lines surround SG cells, or the lumen. Yellow arrowheads point at centrosomes. For (C) one experiment was quantified and 41 embryos were analyzed.

### Supplementary Figure 2

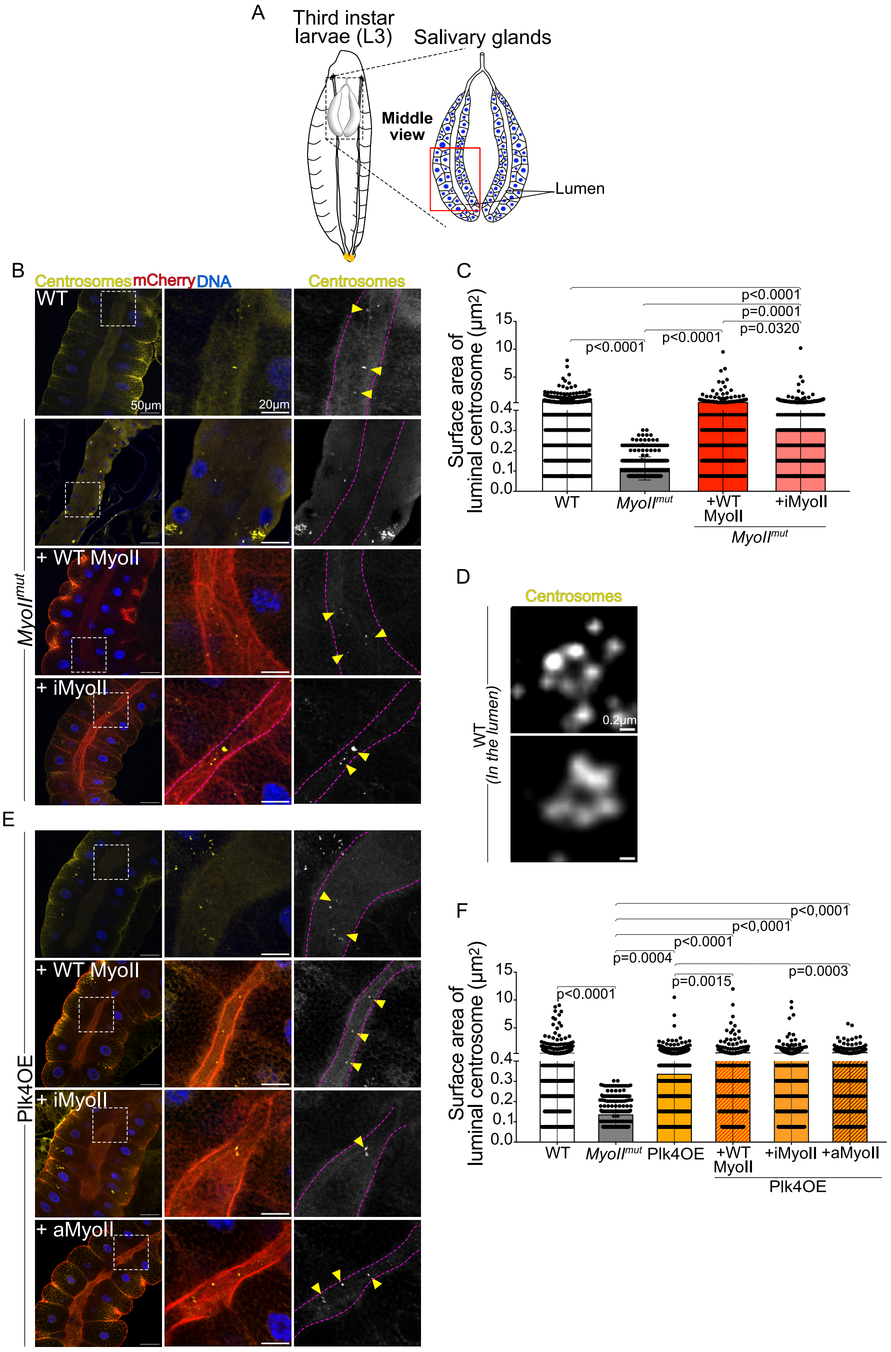

**Supplementary Figure 2- Characterization of centrosome behavior in larva SGs**
(A) Schematic drawing of SGs in L3. On the left an L3 larvae is depicted with anterior positioned at the top. On the right, the SG pair showing a middle view of the polyploid secretory cells with nuclei depicted in blue and the tubular lumen. (B and E) On the left, representative immunostaining images of L3 SGs of the indicated genotypes. Centrosomes in yellow, mCherry-tagged proteins in red and DNA in blue. The white squares mark the luminal regions shown at higher magnification views in the mid panel. On the right, the centrosome channel is shown in grey. Pink dashed lines surround the lumen and yellow arrows point at centrosomes. (C and F) Bar and dot plot graphs showing the individual surface of luminal centrosomes of the indicated genotypes. (D) Representative images obtained by super resolution microscopy of WT luminal centrosomes. iMyoII=inactive MyoII; aMyoII= active MyoII; Statistical significance is shown and determined by ordinary one-way ANOVA tests. Bars show the mean ± SD. For (C), at least three experiments and for (F) four experiments were quantified and 10 SGs were analyzed per condition

### Supplementary Figure 3

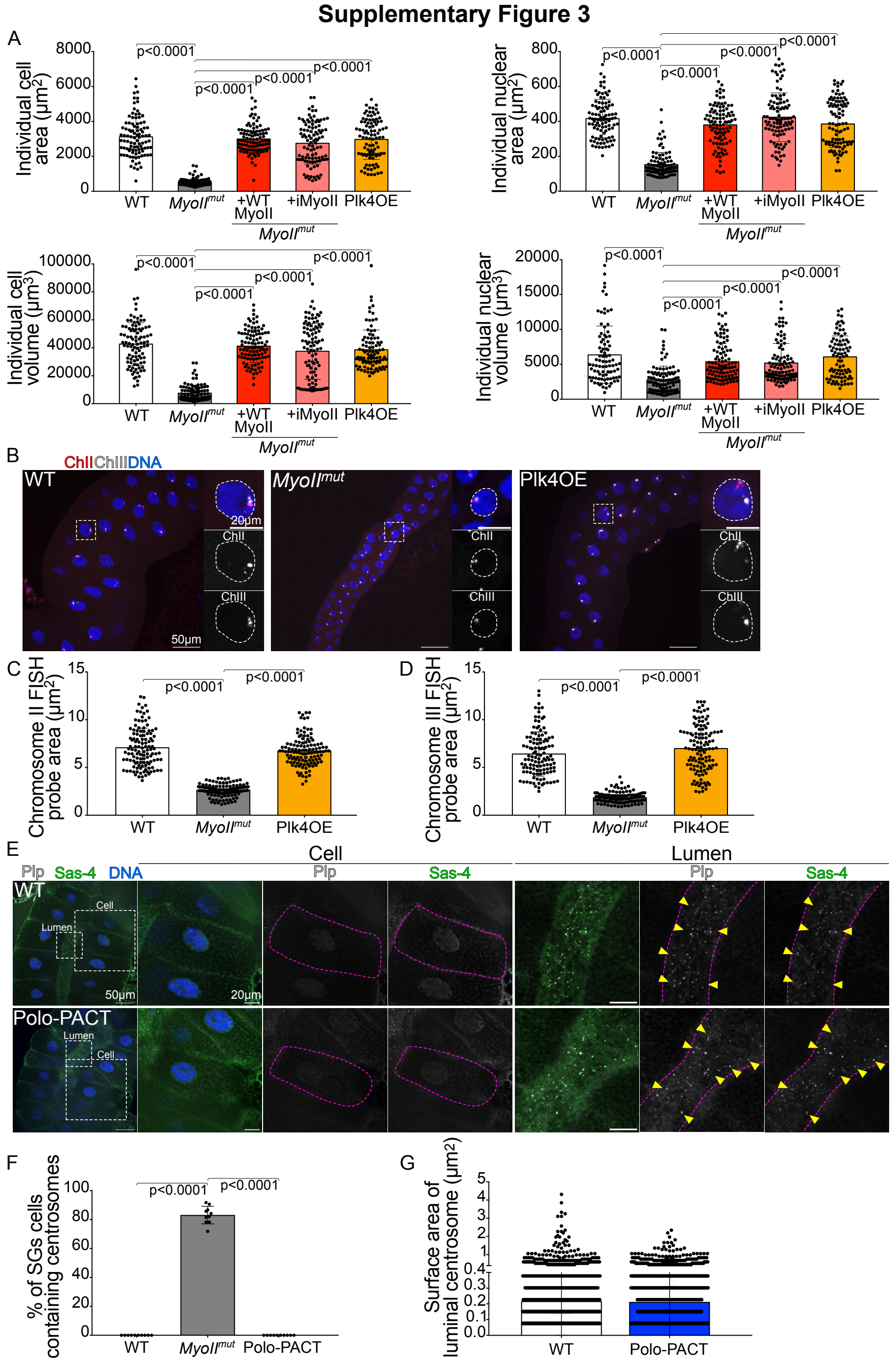

**Supplementary Figure 3- Characterization of *MyoII*$^{mut}$ and Plk4OE SGs**

(A) Bar and dot plot graphs showing individual cell or nuclear area and volume of the indicated genotypes. (B) Representative immunostaining images of L3 SGs of the indicated genotypes showing FISH probes for Chromosome II (red) and Chromosome III (grey). DNA in blue. The white squares mark the cells shown on the right in merged panels or as separated panels in grey and the white dashed circle surround the nuclei. (C-D) Bar and dot plot graphs showing Chromosome II (C) and Chromosome III (D) FISH probe area of the indicated genotypes. (E) Representative immunostaining images of L3 SGs of the indicated genotypes. Centrosomes are shown through the localization of two centrosome markers Plp (grey) and Sas-4 (green). DNA in blue. The white squares mark the two SG regions that are shown at higher magnifications on the right as merged panels or as separate panels showing centrosome markers in grey. These regions correspond to a cell (mid panels) and to the lumen (right panels). The pink dashed lines surround SG cells or the lumen. Yellow arrowheads point at centrosomes. (F) Bar and dot plot graph showing the percentage of SG cells containing centrosomes in the indicated genotypes. (G) Bar and dot plot graph showing the individual surface of luminal centrosomes of the indicated genotypes.
Statistical significance is shown and determined by ordinary one-way ANOVA tests (A, C, D, F) and Mann-Whitney test (G). Bars show the mean ± SD. For (A, C-D), three experiments and for (F-G), two experiments were quantified with a minimum of 10 SGs analyzed per condition.

### Supplementary Figure 4

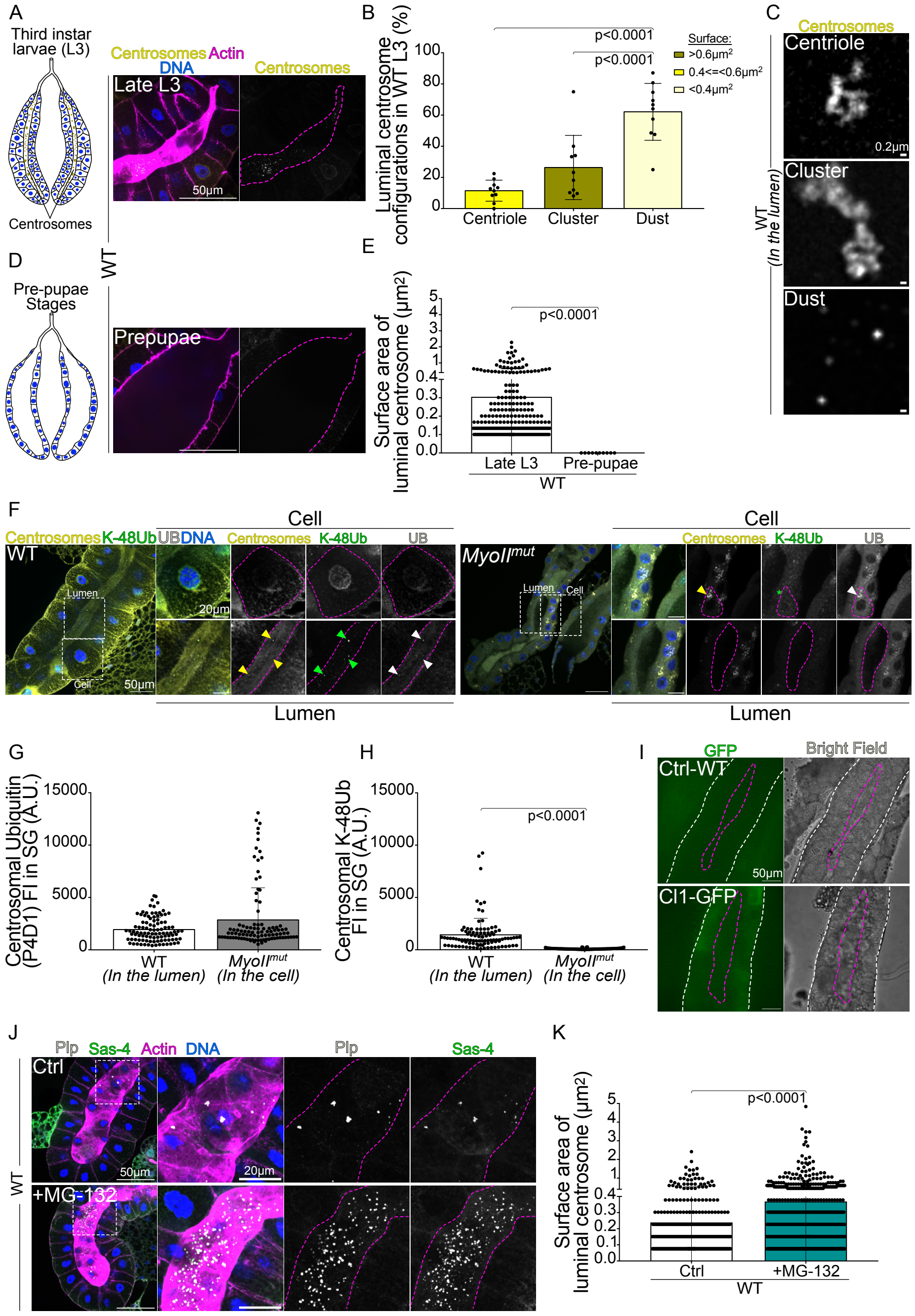

**Supplementary Figure 4- Centrosomes are degraded in the lumen of WT SGs**

(A and D) Schematic drawing of L3 SGs (A) and pre-pupae stages SGs showing centrosomes localizing in a narrow lumen (A) or absent from a wide lumen (D). In both drawings, SGs are depicted with anterior positioned at the top and nuclei in blue. On the right, representative immunostaining images of WT L3 (A) and pre-pupal SGs (D) showing centrosomes in yellow, actin in magenta and DNA in blue in the first panel and centrosomes in the right panel. The pink dashed line marks the lumen. (B) Bar and dot plot graph showing luminal centrosomal configurations in WT L3. (C) Illustrative super resolution images of luminal centrosomes as indicated. (E) Bar and dot plot graph showing individual surface of luminal centrosome in WT at the indicated developmental stages. (F) On the left of each panel, representative immunostaining images of L3 SGs of the indicated genotypes. Centrosomes in yellow, K-48Ub and UB markers shown in green and grey respectively with DNA in blue. The white squares mark two regions shown as insets on the right-hand side. These correspond to cells (top) or luminal regions (bottom). Pink dashed lines surround the cell or the lumen and the colored arrowheads point to the label of the same color. (G-H) Bar and dot plot graphs showing centrosomal P4D1 and K-48Ub FI (A.U.) of the indicated genotypes. (I) Representative images of a SGs expressing the proteasome activity reporter CI1-GFP compared to WT SGs. On the left, the GFP fluorescence is shown andon the right, the corresponding bright field. The white and pink dashed lines surround SGs and the lumen. (J) Representative immunostaining images of WT L3 treated or not with the proteasome inhibitor MG-132, showing two centrosome markers, Plp (grey), Sas-4 (green), actin (magenta) and DNA in blue. The white squares mark the region shown at higher magnifications on the right as merged channels or individual centrosome labeling in grey channels. (K) Bar and dot plot graphs showing individual surface are of luminal centrosome according to the indicated treatment. Statistical significance is shown and determined by ordinary one-way ANOVA tests (B) and Mann-Whitney test (E, G, H and K). Bars show the mean ± SD. For (B) four experiments, (E,G-H) two experiments and for (K) three experiments were quantified with a minimum of 10 SGs analyzed per condition.

#### Supplementary Figure 5

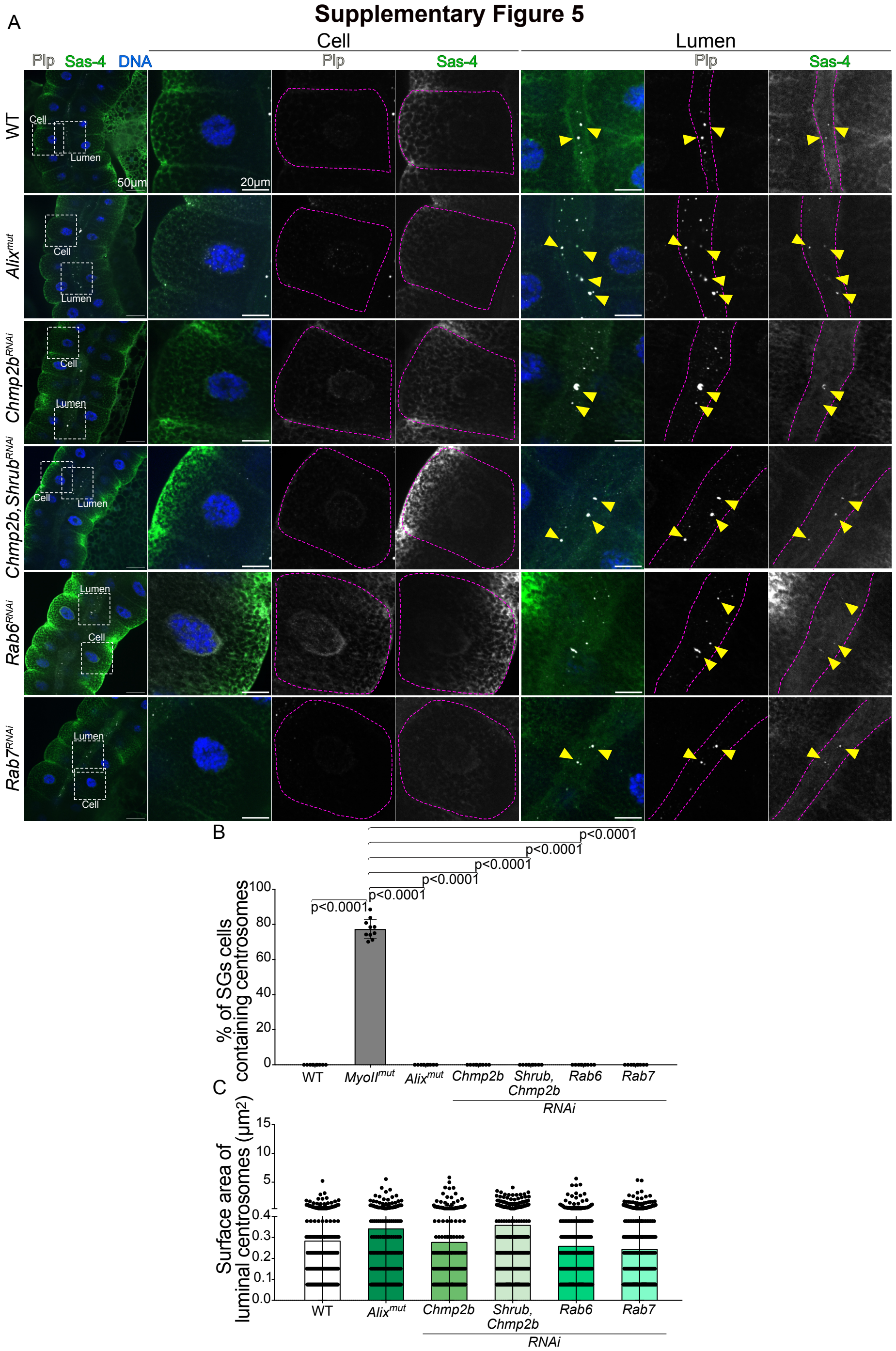

**Supplementary Figure 5- Analysis of centrosomal markers in different genotypes affecting secretion**

(A) On the left, representative low magnification immunostaining images of L3 SGs of the indicated genotypes. Centrosomes are shown as co-localization of two centrosomal markers- Plp (grey) and Sas-4 (green) with DNA in blue. The white squares mark the two SG regions that are shown at higher magnifications on the right as merged panels or as separate panels showing Plp and Sas-4 in grey channels. These regions correspond to a cell (mid panels) and to the lumen (right panels). The dashed pink lines surround the cell or the lumen and the yellow arrowheads point at centrosomes. (B and C) Bar and dot plot graphs showing the percentage of SGs cells containing centrosomes (B) and surface of luminal centrosome (C) of the indicated genotypes. Statistical significance is shown and determined by ordinary one-way ANOVA tests (B-C). Bars show the mean ± SD. For each graph, at least two experiments were quantified, and a minimum of 10 SGs were analyzed per experiment.

**Supplementary Figure 6**

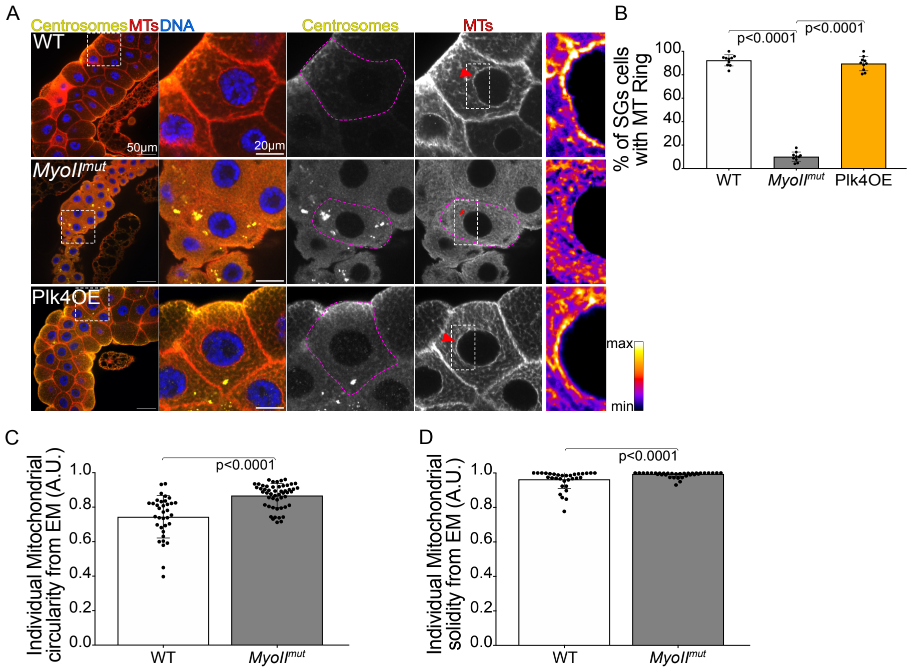


#### Supplementary Figure 6- Analysis of the MT cytoskeleton in SGs

(A) On the left, illustrative low magnification immunostaining images of L3 SGs of the indicated genotypes. Centrosomes in yellow, MTs in red and DNA in blue. The white squares mark a SG cell that is shown at higher magnifications on the right as merged panels or as separate panels showing centrosomes and MTs in grey channels. At the far right, a region displaying fire LUT of MTs is provided. (B) Bar and dot plot graphs showing the percentage of SG cells displaying a MT ring associated with the nuclear envelope in the indicated genotypes. (C and D) Bar and dot plot graphs showing individual mitochondrial circularity (C) and solidity (D) of the indicated genotypes. Statistical significance is shown and determined by ordinary one-way ANOVA test (B) and Mann-Whitney tests (C-D). Bars show the mean ± SD. *For (B*) three experiments and for (C-D) two experiments were quantified, and a minimum of 10 SGs were analyzed (B) and a minimum of 35 mitochondria (C-D).

**Movie 1: Time lapse of embryonic SGs**

Time-lapse movie of embryos (either whole SG or partial SG) at stage 15-17, expressing Sas-4-GFP (yellow) and MyoII-mCherry (red). Yellow arrowhead pointing to the centrosome exiting the cell toward the lumen. Scale bar (10µm for whole SG and 1µm for the crop) and time interval of 30sec.

**Movie 2: 3D reconstruction of WT SG**

3D reconstruction of WT embryonic SG at stage 13-14, showing the full SG (green), the lumen (magenta) and the centrosomes (color coded based on their distance to the lumen as indicated).

**Movie 3: 3D reconstruction of *MyoII$^{mut}$* SG**

3D reconstruction of *MyoII$^{mut}$* embryonic SG at stage 13-14, showing full SG (green), the lumen (magenta) and centrosomes (color coded based on their distance to the lumen as indicated).
